# The effects of human presence, restraint, and stressed neighbors on corticosterone levels in domesticated budgerigars (*Melopsittacus undulatus*)

**DOI:** 10.1101/2025.02.10.637489

**Authors:** Dustin G. Reichard, Kelly V. Summers

## Abstract

Limiting stress during interactions between captive animals and humans is important for effective husbandry. One physiological change during the vertebrate stress response is the release of glucocorticoid hormones. Here, we measured plasma corticosterone in female domesticated budgerigars (*Melopsittacus undulatus*) to test whether human presence, restraint, or removal and return of a newly stressed neighbor increases corticosterone. The presence of humans for 15 minutes at the beginning of the experiment did not significantly elevate corticosterone above baseline levels, suggesting that birds that acclimate to humans are minimally affected by their presence. However, at the end of the experiment after multiple blood sampling events, the same human presence test significantly elevated corticosterone above baseline and human presence levels measured at the beginning of the experiment. Thus, repeated blood sampling could cause progressively stronger stress responses to human presence. Restraint-induced corticosterone levels were significantly higher than all other treatments, indicating that human handling activates the stress response. After stressed birds were returned home, corticosterone levels of their neighbors were significantly higher than baseline at 25- and 45-minutes post-return. However, the 25- and 45-minute corticosterone levels were not significantly different from each other, or levels induced by human presence at the beginning or end of the experiment. This outcome suggests that social transmission of stress was limited across the sampled time frame. These data highlight the importance of evaluating the costs and benefits of different human-animal interactions in captivity, including in domesticated species that are more tolerant of forced proximity to humans.

**Research Highlights:**

- Human presence only elevated plasma corticosterone after multiple blood sampling events.
- Restraint significantly increased corticosterone levels relative to all other treatments.
- Minimal evidence for social transmission of stress.

## Introduction

The stress response is an adaptive process that helps organisms survive and maintain homeostasis despite challenging and unpredictable environments. For example, when a vertebrate encounters a predator, their neuroendocrine system activates a myriad of physiological and behavioral changes to help manage the threat (Sapolsky et al. 2000). Changes such as increased heart rate and blood pressure (Ulrich-Lai and Herman 2009; Vitousek et al. 2010), elevated levels of neurotransmitters and hormones that promote movement and metabolism (Figueiredo et al. 2003; Jones et al. 2016; Lustberg et al. 2022), and a redirection of resources away from nonessential processes like reproduction are beneficial because they increase the likelihood of survival (Wingfield and Sapolsky 2003). However, that same stress response can become costly and maladaptive if it is repeatedly activated in a chronic fashion, such as in the presence of a persistent and unavoidable stressor (Sapolsky et al. 2000; French et al. 2006; DuRant et al. 2016).

Captive animals experience relatively safe and stable environments that are fundamentally different from the natural environments where they evolved and adapted. Both the novelty of captivity and an inability to escape makes it more likely that the stress response will be activated by both threatening and nonthreatening stimuli, potentially leading to chronic stress and many adverse effects (Boonstra 2013). Overactivity of the stress response can cause negative health outcomes such as reduced immunity and weight loss and an increase in problematic behaviors such as aggression and self-harm (Sapolsky et al. 2000; Costa et al. 2016; Fischer and Romero 2019). It can also decrease sexual activity and gamete production (Moore and Jessop 2003; Wingfield and Sapolsky 2003), which can reduce the effectiveness of captive breeding programs (Mason 2010). Thus, identifying and minimizing stressors in the captive environment and evaluating the magnitude of their effects on animal welfare is essential for effective husbandry.

The presence of human caretakers represents a potentially unavoidable stressor for captive animals. Although some individuals can acclimate to the presence of humans over time or be trained to associate humans with positive outcomes, any human interaction involving a novel stimulus or a necessary but invasive procedure such as anesthesia or blood sampling can activate the stress response (Morgan and Tromborg 2007; Fischer and Romero 2019). The activation of the stress response in one individual can also have cascading effects as stress can be socially transmitted (Brandl and Farine 2024). When an individual becomes stressed, mates or group members that recognize a shift in their behavior or other cues may activate their own stress response (Akyazi and Eraslan 2014; Carnevali et al. 2017). A similar effect occurs when an individual is removed from a social group, which disrupts dominance hierarchies and can cause elevated levels of aggression and stress (Sapolsky 1992; Carvalho et al. 2018). Consequently, it is important to not only assess the effects of a human interaction on the focal individual, but also the entire social group.

In this study, we examined the effects of three different potential stressors on the corticosterone levels of female domesticated budgerigars (*Melopsittacus undulatus*). Corticosterone is the primary glucocorticoid hormone released by the adrenal cortex in birds that regulates glucose metabolism and circadian rhythms, among other processes (Sapolsky et al. 2000). Corticosterone levels in the blood plasma typically increase within minutes of exposure to a stressor, making it a commonly used biomarker of the stress response, including in previous studies of budgerigars (Lupu, C. and Robins, S. 2013; Young and Hallford 2013; Medina-García et al. 2017). Budgerigars are highly social psittacines endemic to Australia that were imported to Europe in the 1840s, which is the approximate start of their domestication (Steiner 1939). It is likely that domesticated budgerigars exhibit lower stress reactivity than wild caught budgerigars because domestication involves selectively breeding individuals that respond positively to humans (Fallahsharoudi et al. 2015; Homberger et al. 2015; Karaer et al. 2023), but that hypothesis remains untested in this species.

Over a 14-week period, we collected corticosterone samples bi-weekly so that each budgerigar experienced every treatment and could recover in between blood collection events. We measured corticosterone levels after the following treatments: (1) baseline within 3-minutes of exposure to humans, (2) presence of humans for 15 minutes in the room, (3) 25-minutes of handling and restraint stress by humans, (4) 25 and 45 minutes after the removal and return of a newly stressed neighbor. We predicted that the presence of humans at the beginning of the experiment would not significantly elevate corticosterone above baseline levels because our subjects were exposed to their human caretakers daily and had likely acclimated. However, we predicted that human handling and restraint as well as the removal and return of a stressed neighbor would significantly elevate corticosterone levels due to the severity of these disruptions. Finally, we predicted that at the end of the experiment, birds would elevate corticosterone levels in response to human presence in the room owing to the repeated instances of blood sampling and negative interactions with humans. Understanding the effects of these realistic scenarios will provide insights for minimizing stress during essential husbandry of captive animals.

## Materials and Methods

### Study Subjects

We sampled nine female budgerigars (*Melopsittacus undulatus*) from two color morphs (White: 4, Yellow: 5). These birds were also used in a previous study of diet and immune function that involved repeated blood sampling and was completed two months before the start of the current study (see Eggleston et al. 2019). The only interactions that these birds had with humans in the intervening months were related to daily maintenance (i.e., no blood sampling occurred). Full details of the housing conditions can be found in Eggleston et al. (2019). Briefly, all birds were purchased from a wholesale animal retailer and housed at Ohio Wesleyan University (Delaware, Ohio, USA). Birds were kept in individual cages (38.1 x 48.3 x 50.8 cm), three cages per shelf, and had constant visual and auditory contact with the two neighbors on their shelf. Food and water were available *ad libitum*, and each was replaced daily. Photoperiod was consistently 12 light hours: 12 dark hours and room temperature was maintained at 23°C. This research was reviewed and approved by the Ohio Wesleyan University Institutional Animal Care and Use Committee (Protocol No. 2016-2017-33).

### Treatments and Blood Sampling

We collected a blood sample from each budgerigar every two weeks over a 14-week period, but the conditions surrounding each blood sample varied according to five different treatments (Table 1). Every sample was collected between 16:15-16:45 EST (21:15-21:45 GMT) to control for circadian variation in corticosterone levels and coincide with the bird’s normal daily health checks. We punctured the brachial vein with a 26-gauge needle and collected 50 uL of blood into a heparinized microhematocrit tube. Samples were spun at 10,000 rpm for 10 minutes and the plasma was drawn off and stored at −20 C until the hormone assay (see below). The same three researchers collected all the blood samples, and one of those researchers conducted all of the husbandry.

**Table 1.** Experimental timeline showing the treatment that corresponds to each blood sampling event. Treatments are described in detail in the main text. Each bird was sampled only once each week, which explains the reduced sample size for each treatment in weeks 8, 10, and 12. The “new neighbors” column indicates whether the birds were housed on a shelf with the same or different neighbors relative to the previous week.

| Week | Sampling Treatment | New Neighbors? |
| --- | --- | --- |
| 0 | None | Yes |
| 2 | Human Presence (N=9) | No |
| 4 | Baseline (N=9) | No |
| 6 | Baseline (N=9) | No |
| 8 | Restraint-induced Stress (N=3)<br>Stressed Neighbor 1 (N=3)<br>Stressed Neighbor 2 (N=3) | Yes |
| 10 | Restraint-induced Stress ($N=3$)<br>Stressed Neighbor 1 ($N=3$)<br>Stressed Neighbor 2 ($N=3$) | Yes |
| 12 | Restraint-induced Stress ($N=3$)<br>Stressed Neighbor 1 ($N=3$)<br>Stressed Neighbor 2 ($N=3$) | Yes |
| 14 | Human Presence ($N=9$) | No |

At the beginning of the experiment (week 0), we randomly arranged the birds in individual cages, three cages to a shelf, across three total shelves. After two weeks of acclimation to their shelfmates, three humans entered the room, chatted in normal voices, and observed the birds without approaching their cages for 15 minutes. Then three birds were rapidly captured (one bird/researcher) and a blood sample was taken in less than three minutes to limit the effects of handling on corticosterone levels (Small et al. 2017). After 24 hours had passed, the process was repeated until all birds were sampled.These samples aimed to test the effects of humans in the room that were not directly interacting with the budgerigars on the bird’s corticosterone levels (Treatment 1).

In weeks four and six, we entered the room and rapidly collected blood samples from three birds each day within three minutes of the room’s door opening. These samples assessed the bird’s baseline corticosterone levels (Treatment 2) and were done twice to test whether these levels were consistent through time. After all baseline samples were completed, we shuffled the cages so that each bird received two new shelfmates and allowed them to acclimate for two weeks until week 8. In weeks eight, ten, and twelve, we needed to enter and exit the room four separate times in a single day to accomplish our blood sampling treatments (see below). To minimize any effect of repeated human presence on corticosterone levels, we turned off the lights before entering the room and turned them back on as soon as we left so that the birds would not see a human approach their cage. Once in the dark room, we rapidly collected one cage from each shelf and moved the cages to an adjacent room to minimize disrupting the remaining birds. This entire process was quick and took less than one minute.

The first cages that we removed were always in the middle of the shelf, which allowed us to later assess the responses of both neighbors on either side of the stressed bird. After the first three caged birds were moved to the new room, we immediately captured each bird and placed it into a small paper bag to simulate restraint-induced stress (Treatment 3). After 25 minutes had passed since we initially picked up the bird’s cages, we removed the birds from their bags and collected a blood sample in less than 3 minutes (∼28 minutes after the onset of the stressor). Previous work in domesticated budgerigars by Medina-García and colleagues (2017) has shown that corticosterone levels peak after 30 minutes of restraint stress. Then these birds were immediately returned to their cages, and the cages were returned to their home shelves.

Twenty-five minutes after the stressed birds were returned to their home shelves, we collected one of the two remaining cages from each shelf and moved them to the adjacent room. Then we caught and collected a blood sample from each bird within 3 minutes of initially picking up their cages. These samples measured the effects of spending 25 minutes with a neighbor that experienced restraint-induced stress (Treatment 4). Forty-five minutes after the initial stressed bird was returned home, we repeated our sampling procedure with the remaining three birds. These samples measured the effects of spending 45 minutes with a stressed neighbor (Treatment 5). The day after these samples were collected, we shuffled the cage locations in a random, but stratified manner so that each bird received two new neighbors for the subsequent sampling. The role played by each bird (i.e., restraint stress, stressed neighbor time 1 or 2) also differed during each sampling so that each bird played each role. The bird receiving the restraint-induced stress treatment was always located in the middle of the shelf. During week fourteen, we repeated the human presence treatment from week 2 to test for any cumulative effects of repeated blood sampling on the bird’s response to humans in their room. Over the entire experiment, we failed to collect a sufficient blood sample within the three minutes at four different time points (restraint stress, *N* = 1; Stressed Neighbor 2, *N* = 1; Human Presence 2, *N* =2), which reduced our sample size for those treatments.

### Hormone Assays

A previous study by Lupu and Robins (2013) measured corticosterone in samples of unextracted budgerigar plasma using an enzyme immunoassay kit (EIA) from Cayman Chemical (Kit No. 500651, Ann Arbor, Michigan, USA), but they did not report the results of an assay optimization. We used an updated Cayman corticosterone EIA kit (Kit No. 501320), but we were unable to achieve parallelism between an eight-point standard curve created from kit reagents and an equivalent displacement curve of pooled plasma samples from the birds used in this study in two attempts (Plate 1: *F*_7_ = 5.03, *P*<0.05, *r*^2^=0.92; Plate 2: *F*_7_ = 8.30, *P*=0.01, *r*^2^=0.61; see below for analysis details). However, we achieved strong parallelism between the kit standard and pooled plasma after extracting the plasma twice with ethyl ether (Plate 3: *F*_7_ = 2.15, *P*=0.33, *r*^2^=0.98).

For the extraction, we added 1 mL of ethyl ether (anhydrous, ≥99% purity, Certified ACS; Thermo Fisher Chemical, Waltham, Massachusetts, USA) to a tube containing the pooled plasma, vortexed, and allowed the sample to separate at room temperature for 20 minutes. Then, we snap froze the sample in a bath of methanol and dry ice, and the supernatant was poured into a new tube and placed in a 40 C heating block to evaporate the ethyl ether. We repeated the process with the remainder of the original sample and added the supernatant to the same tube in the heating block. Once all the ethyl ether had evaporated, we reconstituted the sample in assay buffer from the EIA kit and conducted the assay.

To analyze our samples, we extracted 15 uL of plasma from each sample and reconstituted in 200 mL of assay buffer as described above and followed the manufacturer’s instructions to measure corticosterone. Samples were randomized and divided across two plates, but all the samples from each individual bird were run on the same plate. We used a four-parameter logistic standard curve to calculate corticosterone concentrations (BioTek Gen5 Software version 2.09; Agilent, Santa Clara, California, USA). A pooled plasma extract was shared across plates to measure inter- and intra-plate variation. Interplate variation was 8.0%, but due to a pipetting error, we were only able to measure intraplate variation on one of the two plates (1.7%).

### Statistical Analysis

All statistical tests were conducted in R version 4.4.2 (R Core Team 2024) using the “stats” package unless otherwise noted. Assay parallelism was assessed using an *F*-test to compare the slopes of the standard curve and the displacement curve of pooled plasma or plasma extract for each optimization plate. We used linear regressions to plot the curves against one another and calculate an *r*^2^ value.

Four of the baseline plasma corticosterone samples, including both samples from one individual, were not detectable by our assay, but all the other samples fell on the standard curve. The reported sensitivity of the assay kit is 0.021 ng/mL, and we have no reason to suspect that these non-detectable samples were the result of errors during the extraction or assay procedure. Furthermore, a previous study of domesticated budgerigars also found multiple individuals with undetectable baseline corticosterone samples using a different EIA kit (Medina-García et al. 2017). To maximize our sample size, we included these non-detectable samples as zeroes for analysis. Importantly, removing these zeros only affected the statistical significance of one pairwise comparison (baseline v. 45 minutes with stressed neighbor, *P*=0.088).

We used a paired t-test to determine whether corticosterone levels differed between the two baseline samples. Then we used a linear mixed-effects model (package: “lme4”) to examine differences in corticosterone levels with a modified Bonferroni correction to assess post-hoc pairwise comparisons between treatments (Benjamini and Hochberg 1995; package: “grafify”). In our model, plasma corticosterone was the dependent variable, treatment was the fixed effect, and subject ID was included as a random effect to account for repeated sampling. The residuals of our initial model failed a Shapiro-Wilk test of normality and marginally passed Levene’s test of homoscedasticity, two assumptions of linear mixed models. We resolved these issues by square root transforming the corticosterone data and report the results of that model here.

## Results

Baseline plasma corticosterone levels did not differ significantly between measurements (*t_8_* = −0.60; *P* = 0.56) and were averaged for the remaining analyses. The presence of humans for 15 minutes at the beginning of the experiment did not significantly elevate plasma corticosterone levels above average baseline levels (Table 2, Figure 1). However, the presence of humans at the end of the experiment significantly elevated corticosterone levels both above the average baseline and the presence of humans at the beginning of the experiment (Table 2, Figure 1). Restraint-induced stress significantly increased corticosterone levels relative to all other treatments (Table 2, Figure 1).

**Table 2.** Each cell reports the t-statistic (top) and adjusted P-value (bottom) from pairwise post-hoc comparisons derived from a linear mixed model with a modified Bonferroni correction (Benjamini and Hochberg 1995). Bold values represent significant differences (P<0.05).

|  | Average<br>Baseline<br>(*N*=9) | Human<br>Presence - 1<br>(*N*=9) | Stressed<br>Neighbor -<br>25<br>(*N*=9) | Stressed<br>Neighbor -<br>45<br>(*N*=8) | Restraint<br>Stress<br>(*N*=8) |
| --- | --- | --- | --- | --- | --- |
| Human<br>Presence - 1<br>(*N*=9) | -1.60<br>0.15 | -- | -- | -- | -- |
| Stressed<br>Neighbor - 25<br>(*N*=9) | **-3.33**<br>**0.004** | 1.73<br>0.13 | -- | -- | -- |
| Stressed<br>Neighbor - 45<br>(*N*=8) | **-2.58**<br>**0.02** | 1.03<br>0.33 | 0.64<br>0.53 | -- | -- |
| Restraint<br>Stress<br>(*N*=8) | **-9.17**<br>**<0.0001** | **-7.62**<br>**<0.0001** | **-5.95**<br>**<0.0001** | **-6.38**<br>**<0.0001** | -- |
| Human<br>Presence - 2<br>(*N*=7) | **-4.41**<br>**0.0003** | **-2.91**<br>**0.01** | -1.31<br>0.23 | -1.87<br>0.10 | **-4.30**<br>**0.0003** |

**Figure 1.**
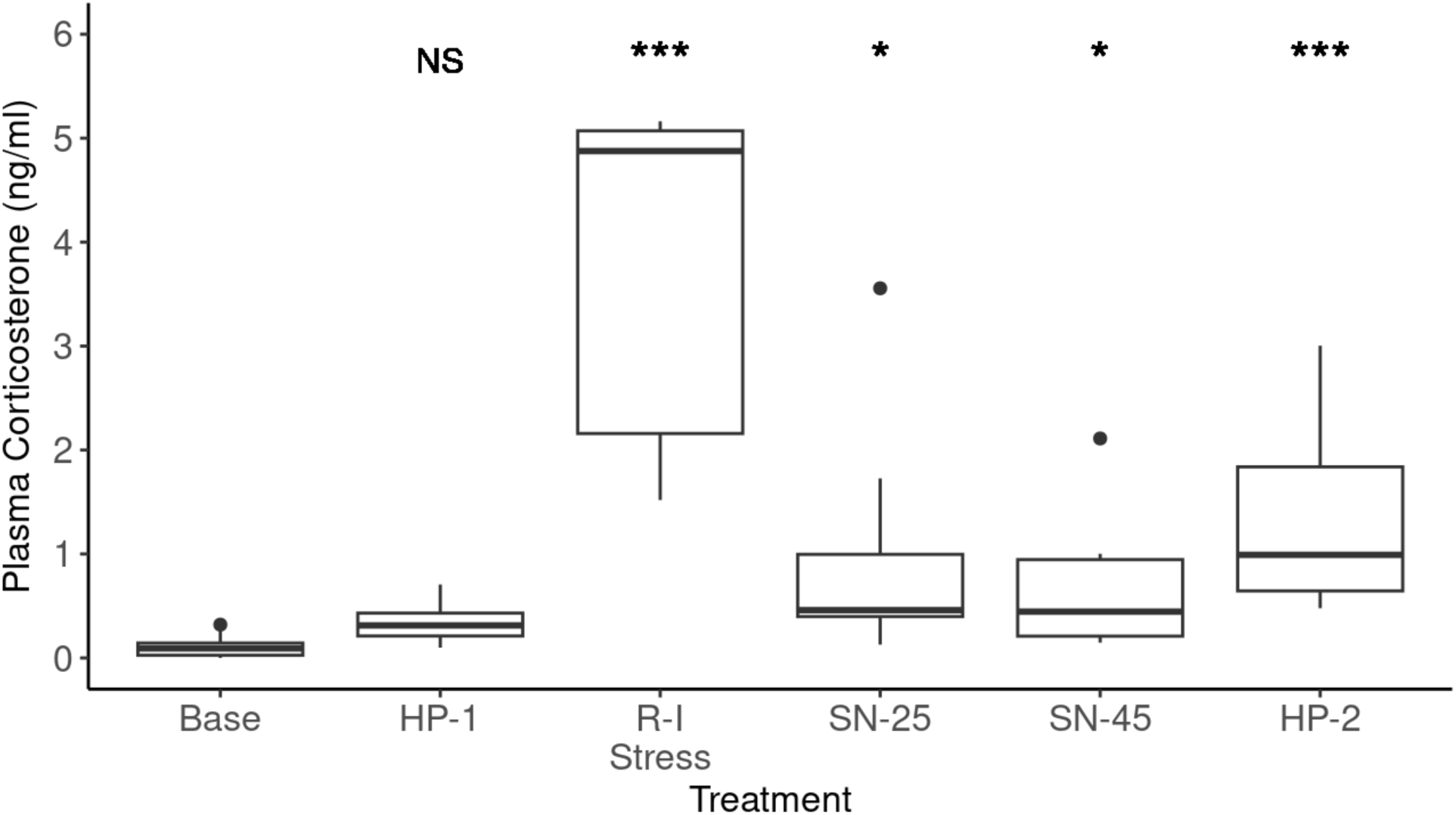
Plasma corticosterone levels in domesticated budgerigars after six different treatments. Abbreviations correspond to blood samples collected with 3 minutes or less of handling under the following conditions: “Base” = less than 3 minutes of human presence and handling (N = 9), “HP” = humans present in birds room for 15 minutes before sample (1 = pre-experiment, N = 9; 2 = post-experiment, N = 7), “R-I Stress” = 25 minutes of restraint in a paper bag (N = 8), “SN” = stressed neighbor returned to neighboring cage before sample (25 = 25 minutes with stressed neighbor, N = 9; 45 = 45 minutes with stressed neighbor, N = 8). Treatments are arranged in chronological order of sampling from left to right along the x-axis. Boxes enclose interquartile range and thick line identifies median. Whiskers enclose 1.5 times the interquartile range and dots represent points that exceed that range. Significance markers represent results of pairwise comparisons between treatment and baseline corticosterone levels only. NS = not significant. ***P<0.001. *P<0.05.

Birds that observed the removal and return of a newly stressed neighbor had significantly higher levels of corticosterone than average baseline levels at both 25 and 45 minutes after the neighbor was returned (Table 2, Figure 1). However, corticosterone levels were not significantly different between birds that spent 25 versus 45 minutes with the stressed neighbor nor did they differ detectably between birds that spent time with stressed neighbors and birds that experienced human presence for 15 minutes at the beginning or end of the experiment (Table 2, Figure 1).

## Discussion

We measured plasma corticosterone, a biomarker of the stress response, in domesticated budgerigars that interacted with humans in a variety of contexts relevant to husbandry and welfare. As predicted, direct handling and restraint for 25 minutes resulted in significantly higher levels of corticosterone than all other treatments. All birds in this study were from a domesticated lineage and had previous experience with blood sampling (Eggleston et al. 2019); however, that past artificial selection and opportunity for acclimation did not eliminate the stress response during a hands-on interaction with humans. Similar results have been observed in other domesticated animals such as chickens (*Gallus gallus*) and silver foxes (*Vulpes vulpes*), that exhibit lower baseline and stress-induced levels of glucocorticoids but nonetheless elevate these hormones in response to human handling and restraint (Trut et al. 2009; Fallahsharoudi et al. 2015). The activation of the stress response and release of glucocorticoids during handling is widely documented among captive animals, and it is relevant in clinical settings as it can affect the results of diagnostic measures such as body temperature, respiratory rate, and white blood cell counts (Greenacre and Lusby 2004; McRee et al. 2018).

The presence of humans in the room for 15 minutes did not significantly elevate corticosterone above baseline levels at the beginning of the experiment. This result is consistent with the birds having acclimated to humans, which entered the room daily for visual wellness checks and replacement of food and water. Consistent, positive interactions with humans can lead to reduced levels of corticosterone during human handling in broiler chickens (Hemsworth et al. 1994), and observations from both livestock and zoos have found similar reductions in fear or stress-related behaviors over time for some species (Hosey 2000; Hemsworth 2003; Sherwen and Hemsworth 2019). It is important to acknowledge that the absence of an increase in corticosterone in this context does not necessarily indicate the lack of a stress response. Glucocorticoid levels do not always elevate in response to stress, such as during certain life history stages like molt in birds, and there are multiple other physiological parameters not measured here, such as heart rate and blood pressure, that could have changed because of the stress response (MacDougall-Shackleton et al. 2019).

At the end of the experiment, after experiencing six separate blood sampling events, budgerigars significantly elevated corticosterone in response to human presence for 15 minutes relative to both baseline levels and the response to human presence at the beginning of the experiment. This outcome suggests that repeated blood sampling may cause birds to develop an anticipatory activation of the stress response in the presence of humans, which could be harmful to health and welfare if these reactions are occurring every time a human enters the room (Morgan and Tromborg 2007). This result emphasizes the importance of minimizing the number and severity of negative interactions with humans over time. However, in some cases, animals are capable of acclimating to repeated negative interactions with humans when the outcome is always survival with minimal injuries (Romero 2004). If budgerigars are capable of acclimating to repeated blood sampling, our data suggest that either more sampling events or changes to the sampling methodology (e.g., Gouveia and Hurst 2019) are necessary to reach that goal.

The removal of animals from social groups is sometimes required for clinical or experimental procedures, and any stress incurred by that animal could be transmitted socially to its groupmates when it returns (Akyazi and Eraslan 2014; Carnevali et al. 2017). We found that the removal and return of a newly stressed neighbor significantly elevated corticosterone levels above baseline for birds in neighboring cages after 25 and 45 minutes. However, corticosterone levels in neighbors did not differ detectably from levels observed after spending 15 minutes in the presence of humans at both the beginning and end of the experiment. Collectively, these results suggest that although neighbors of the stressed bird experienced their own stress response, it was more likely caused by human presence in the room than social transmission of stress. Had social transmission occurred, we would have expected corticosterone levels to be significantly higher than both baseline and human presence levels. The lack of social transmission of stress could have been caused by the birds being individually caged, which made physical contact, such as allopreening, impossible and limited affiliative behavior (Morales Picard et al. 2020). In addition, evidence from zebra finches (*Taeniopygia guttata*) has suggested that the effects of social disruption could be transient as birds exhibited higher levels of corticosterone 10 minutes after the removal of a neighbor or partner, but not after 30 minutes (Banerjee and Adkins-Regan 2011). Thus, our sampling may have occurred too late to observe any effects of the social disruption on corticosterone levels. Nevertheless, this result suggests that social transmission of stress has minimal effects on the stress response of budgerigars housed in individual cages.

Corticosterone levels in captive parrots tend to be lower at baseline (Vidal et al. 2019) and return to baseline faster after activation of the stress response (Cabezas et al. 2013) than levels observed in wild parrots. The corticosterone levels observed in our domesticated budgerigars are lower than the levels measured in two other published measurements of plasma corticosterone from captive birds of this species (Lupu, C. and Robins, S. 2013; Medina-García et al. 2017). In the most comparable study, Medina-García and colleagues (2017) also found undetectable baseline samples of corticosterone, but their average restraint-induced value was more than triple what we observed (<u>*X*</u> = 3.83 ng/mL v. 11.62 ng/mL). However, the length of the restraint was longer (25 v. 30 minutes), subjects were male rather than female, housing was in large flight cages rather than individual cages, and all birds were sourced from a different breeder than this study, which could partially explain these differences. Additionally, we only completed a double ethyl ether extraction rather than a triple extraction, which likely resulted in marginally lower corticosterone measures as we were unable to recover all the hormone in each sample.

In conclusion, the results of this study confirm that handling and restraint activates the stress response even in a species that has been domesticated and selectively bred for close relationships with humans. Furthermore, when budgerigars are handled and experience negative interactions like blood sampling repeatedly, they seem to develop an anticipatory stress response activated by the presence of humans in the same room. These outcomes highlight the importance of minimizing the occurrence of human handling and restraint, and future work should explore methods for minimizing the stress response when handling cannot be avoided. Examining the effects of different restraint methods, blood sampling locations (e.g., brachial v. jugular), and approaches to acquiring blood have yielded promising results in other species and should be tested in psittacines (Kannan and Mench 1996; Arnold et al. 2008; Tsai et al. 2015). Finally, we found limited evidence for social transmission of stress in individually caged budgerigars with only visual and auditory access to neighbors. Future work should address whether social transmission of stress is more common in budgerigar flocks held in larger flight cages where direct physical interactions are possible.

## Data Availability Statement

All data and code related to this study are available at Mendeley Data, DOI: 10.17632/pdn9kb984r.1.

## Funding Statement

This study was funded by Ohio Wesleyan University.

## Conflict of Interest Statement

The authors have no conflicts of interest to declare.

## Ethics Statement

This research was reviewed and approved by the Ohio Wesleyan University Institutional Animal Care and Use Committee (Protocol No. 2016-2017-33).

## Notes

https://doi.org/10.17632/pdn9kb984r.1

### Competing Interest Statement

The authors have declared no competing interest.

